# Chromosome-level genome assembly of the photobiont alga *Trebouxia* sp. ‘A48’ from *Xanthoria parietina* provides new insight into the lichen symbiosis

**DOI:** 10.1101/2025.05.01.651714

**Authors:** Gulnara Tagirdzhanova, Jasper Raistrick, Nicholas J. Talbot

## Abstract

- Lichens are symbiotic assemblies consisting of multiple organisms, chiefly a fungus and a photosynthetic microorganism or the photobiont. Among diverse photobionts, the most prevalent is the chlorophyte alga *Trebouxia*.
- We produced a chromosome-level assembly of *Trebouxia* sp. ‘A48’, a photobiont of *Xanthoria parietina*. The genome was assembled into 20 contigs, of which 16 had telomeric repeats on both ends and likely represent complete chromosomes. We compared the genome to genomes of other *Trebouxia* species and analyzed it to investigate the biology of the species, including its lifecycle and adaptations to lichen lifestyle.
- *Trebouxia* sp. ‘A48’ genome is haploid and encodes genes involved in sexual reproduction and meiosis. The predicted secretome is enriched in hydrolases and redox enzymes and contains carbohydrate-binding proteins potentially involved in cell-to-cell recognition and adhesion. We identified genes potentially involved in carbon concentrating and confirmed two instances of ancient horizontal gene transfer from fungi.
- The genome and the strain of *Trebouxia* sp. ‘A48’ are available for the community for research on algal evolution and lichen symbiosis.

## Introduction

The textbook example of symbiosis, the lichen symbiosis, is centered around the relationship between a fungal partner called the mycobiont, and a photosynthetic symbiont, known as the photobiont (Spribille *et al*., 2022). Lichen photobionts are diverse and include both prokaryotic cyanobacteria and eukaryotic microalgae, and among both groups the lichen lifestyle has emerged several times independently (Scharnagl *et al*., 2023). Our understanding of the mechanics and evolution of the lichen symbiosis relies on generating sufficient information from both key symbionts. However, compared to their fungal partners, lichen photobionts have received little attention. This gap has begun to close recently with new studies investigating the genomics and evolution of chlorophyte photobionts (Puginier *et al*., 2024; Poquita-Du *et al*., 2024; Gazquez *et al*., 2024). However, only a handful of high-quality genomes of lichen algae are currently available, because most existing lichen photobiont genome sequences originate from short-read, often metagenomic, data and tend to be highly fragmented.

The Chlorophyta alga *Trebouxia* (Order Trebouxiales, Class Trebouxiophyceae) is the most common lichen photobiont, estimated to participate in nearly half of all described lichen symbioses (Sanders & Masumoto, 2021). *Trebouxia* algae are diverse, and while only around 30 species have been formally described, the genus currently includes 113 candidate species split into four clades: clade A (*T. arboricola/gigantea* group), clade C (*T. corticola/galapagensis/usneae* group), clade I *(T. impressa/gelatinosa* group), and clade S (*T. implex/letharii/jamesii* group) (Muggia *et al*., 2020). In spite of the importance of *Trebouxia* as a photobiont, our understanding of its biology has been severely limited by the absence of high quality genome sequence information.

In this study, we present the genome sequence of a *Trebouxia* photobiont of the common sunburst lichen *Xanthoria parietina* (Figure 1), which has long served as a model system for studies of lichen development, physiology, and genetics (Korhonen & Kallio, 1987; Honegger, 1996; Honegger *et al*., 2004). *X. parietina* lichens involve several *Trebouxia* lineages, with the majority coming from the *T. decolorans* species complex (Nyati *et al*., 2014). We have generated a chromosome-level assembly of the photobiont genome and used this to investigate the life cycle, symbiotic lifestyle, and evolutionary history of the symbiotic alga.

**Figure 1.**
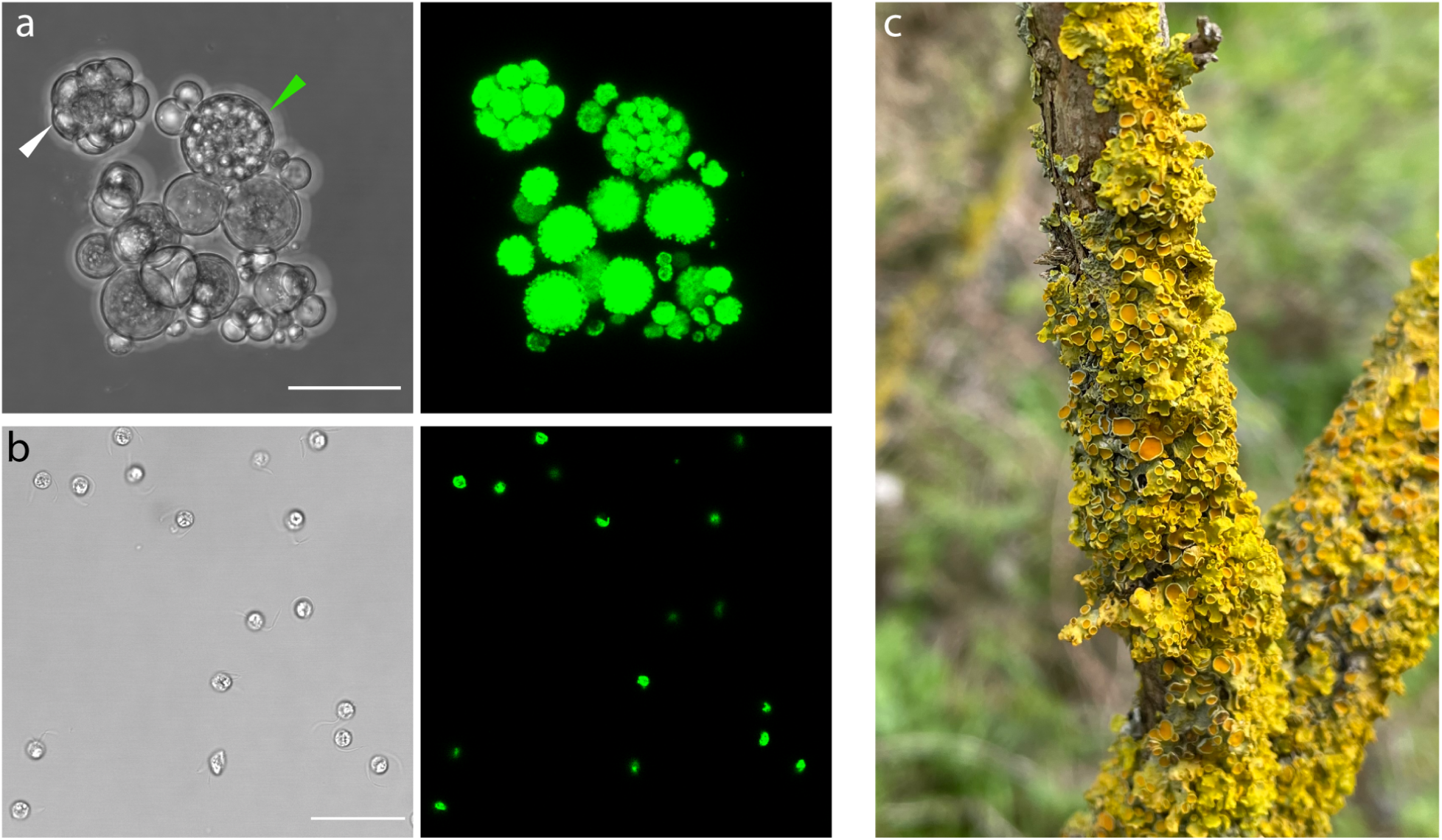
*Trebouxia* sp. ‘A48’. A-B. Micrographs of *T.* sp. ‘A48’ cultured from a *X. parietina* thallus. The left panel shows bright-field microscopy; the right panel shows chlorophyll autofluorescence. Scale bar = 25 μm. **A.** A cluster containing mature and young *T.* sp. ‘A48’ cells, as well as an autosporangium (white arrow) and a zoosporangium (green arrow). **B.** Flagellated cells of *T.* sp. ‘A48’. **C.** A thallus of *X. parietina*.

## Methods

### Strain isolation

The algal strain was isolated from a thallus of *X. parietina* lichen collected from an oak branch in Norwich Research Park (Norwich, UK; 52.622991°N, 1.221588°E). We followed the isolation procedure of Yoshimura *et al*. (2002): a thallus fragment was washed under a jet of water, homogenized, and plated on solid Bold’s Mineral Medium. Plates were incubated on a bench top for one week and then moved to a growth chamber, where they were incubated at 18°C with a 12 hour light/dark cycle at 550 µmol m^−2^ s^−1^ light levels. After six weeks of incubation, several algal colonies were re-plated on solid Trebouxia Organic Nutrient Medium (TONM) (Yoshimura *et al*., 2002) with rifampicin (the final concentration of 50 μg/mL) and carbendazim (40 μg/mL). A single colony from one of the contamination-free plates was then selected and re-plated again. To bulk the culture, we used liquid TONM medium with rifampicin (50 μg/mL) and carbendazim (25 μg/mL) incubated in the conditions described above. To confirm the axenic status of the culture, we cultured it on solid lysogeny broth (LB) medium and incubated it for two days at 28°C and 37°C. The algal strain was deposited to The Culture Collection of Algae at the University of Göttingen, Germany (SAG) as NA 2025.008 (temporary ID, permanent ID is pending).

### Nucleic acid extraction and sequencing

We concentrated 100 mL of liquid algal culture by centrifugation at 4,000 rpm for 5 min. The cell pellet was snap-frozen and freeze-dried. We homogenized approximately 80 mg of dried material using Geno/grinder at 1,300 rpm for 1 min. Genomic DNA was extracted using a NucleoBond High Molecular Weight DNA Kit and purified using a Qiagen genomic Tip20 kit and a Circulomics Short read eliminator kit with a 25-kb cut-off. A sequencing library was prepared with a Native Barcoding Kit 24 V14 and sequenced on a PromethION P2 Solo (P2S) device with a PromethION Flow Cell FLO-PRO114M to 22.4 Gbp of raw data. Base calling was performed with Dorado v7.4.13. Genomic sequencing and assembly were carried out by Future Genomics (Leiden, Netherlands).

For total RNA extraction, we harvested and homogenized algal material as described above. Total RNA was extracted with a RNEasy Plant Mini Kit; the mRNA library was prepared and sequenced by Novogene on an Illumina machine to 8 Gbp of PE150 data.

### Genome assembly

Draft assembly was performed with Hifiasm v0.24.0 (Cheng *et al*., 2021). We used samtools v1.12 (Li *et al*., 2009), Minimap2 v2.24-41122 (Li, 2018), MetaBAT v2.15 (Kang *et al*., 2019), and blast+-2.9.0 (Camacho *et al*., 2009) to examine the contigs, including their coverage, GC content, and BLAST matches to the NCBI nt database (Figure S1a). For the nuclear genome, we retained contigs with GC content 48-50% and coverage 180-210x. One additional contig with coverage 390x was identified as hybrid, where a fragment of the nuclear and plastid genome were fused. By examining the read mapping file, we identified the misassembly point (Figure S1b) and manually split the contigs. To identify other cases of misassembly resulting from telomeric repeats, we identified and examined every instance where telomeric repeats were present more than 500 bp away from the contig terminus. Several similar misassembly points were thus identified and removed (Figure S1c). Notably, one of the contigs included in the final genome assembly had a majority of BLASTn hits to animal genomes (Figure S1a). We considered these hits erroneous and included the contig in the assembly for following reasons: (a) the contig is over 2 Mbp long with no plausible misassembly sites and has the same coverage depth as the rest of the nuclear genome; (b) all but one hits to animal genomes originate from a fragment only 1,300 bp long; (c) all these hits were to fish genomes, which can plausibly be contaminated with algal sequences; (d) high-quality hits to lichen-associated algae can be obtained from the same region. We confirmed the quality of the nuclear genome assembly using BUSCO v4.0.6 (Seppey *et al*., 2019) with the chlorophyta_odb10 database. To align the nuclear genome assembly to the genome of *Trebouxia* sp. SL0000003 (PRJEB86095), we used Minimap2 v2.24-41122 (Li, 2018) with the -x asm20 flag. The synteny plot was generated with syntenyPlotteR (Quigley *et al*., 2023).

As the mitochondrial genome failed to assemble as a single contig, we reassembled it from reads identified as mitochondrial. First, we extracted all reads that mapped to the group of contigs with BLAST matches to Chlorophyta mitochondria (Figure S1a). We then selected the top 30,000 reads and re-assembled them using Hifiasm v0.18.5 (Cheng *et al*., 2021).

### Genome annotation

We masked repeats using RepeatMasker v4.0.9 (https://www.repeatmasker.org/) with a custom repeat library prepared with RepeatModeler v2.0.3 (Hua-Van & Capy, 2024) from the genomic assembly. The repeat-masked genome was then used for gene prediction using the funannotate pipeline v1.8.15 (Palmer & Stajich, 2020). First, we used the funannotate ‘train’ module to train Augustus prediction using transcriptomic data. Next, we run gene prediction using Genemark-ES v4.62 (Lomsadze *et al*., 2005) and the funannotate ‘predict’ module, which collected predictions from Augustus v3.3.2 (Stanke & Waack, 2003), GlimmerHMM v3.0.4 (Majoros *et al*., 2004), and SNAP 2006-07-28 (Korf, 2004). We created consensus models with EVidence Modeler v1.1.1 (Haas *et al*., 2008) and annotated tRNA with tRNAscan-SE v2.0.9 (Chan & Lowe, 2019).

The predictions were refined using transcriptomic data with the funannotate ‘update’ module. For functional annotations, we used InterProScan v5.52-86.0 (Paysan-Lafosse *et al*., 2023) and the funannotate ‘annotate’ module, which collected predictions from PFAM v35.0 (Mistry *et al*., 2021), UniProtDB v2023_01(UniProt Consortium, 2023), MEROPS v12.0 (Rawlings *et al*., 2014), dbCAN v11.0 (Yin *et al*., 2012), and BUSCO chlorophyta_odb10 (Seppey *et al*., 2019). We also annotated KEGG ontology using KAAS webserver (Moriya *et al*., 2007). To detect telomeric repeats, we used the script from Hiltunen *et al*. (2021), with “CCCTAAA” as a query. *T.* sp. SL0000003 genome was annotated using the same protocol, with two modifications: due to the lack of RNAseq data, we skipped the ‘train’ and ‘update’ modules; instead, we supplied the predicted proteome of *T.* sp. ‘A48’ as evidence for the ‘predict’ module. To identify genes involved in meiosis and syngamy, we relied on the InterProScan annotations or, in the case of *HOP1*, *HOP2*, and *MER3*, we used genes from *Coccomyxa subellipsoidea* (Fučíková *et al*., 2015) as BLAST queries. To search for gamete-specific genes *FUS1* and *MTD1*, we used sequences from *Chlamydomonas reinhardtii* (AAL14635.1 and AAC49416.1). We searched both the predicted proteome and the original pre-decontamination assembly.

To characterize ploidy, we followed the procedure of Ament-Velásquez *et al*. (2021). We aligned the long read data on the nuclear genome assembly using Minimap2 v2.24-41122 (Li, 2018) and removed duplicated reads with Picard v2.21.2 (https://broadinstitute.github.io/picard/). We called variants using VarScan v2.3.9 (Koboldt *et al*., 2012) with the flags --p-value 0.1 --min-var-freq 0.005. The results were processed using vcfR library v1.15.0 (Knaus & Grünwald, 2017); only positions with coverage depth 200-240x were considered.

For secretome prediction, we used WolfPSORT (Horton *et al*., 2007), deepTMHMM (Hallgren *et al*., 2022), and SignalP v5 (Almagro Armenteros *et al*., 2019). We considered protein as secreted if it has signal peptide identified by SignalP, no transmembrane domains identified by deepTMHMM, and the highest probability of being secreted according to WolfPSORT. We used ClusterProfiler v4.2.2 (Yu *et al*., 2012) for enrichment analysis.

We annotated the plastid and mitochondrial genomes using MFannot (Lang *et al*., 2023) and GeSeq (Tillich *et al*., 2017). To cross-reference the annotations with the transcriptome data, we aligned RNA data onto the organelle genomes using STAR v2.5.4b (Dobin *et al*., 2013). The final annotations were curated manually and visualized with OGDRAW (Greiner *et al*., 2019).

### Phylogenetic analysis

To provide a taxonomic assignment for the genome, we assembled a reference set of *Trebouxia* internal transcribed spacer (ITS) sequences following Muggia *et al*. (2020). For each of the operational taxonomic units (OTUs), we selected at most five representatives for which an ITS sequence was available and added *Myrmecia* sp. 1 IM-2024 (PQ154957.1) as an outgroup (Table S1). To identify ITS sequences within our genome assembly and *Trebouxia* sp. SL0000003, we used a BLASTn search with *Trebouxia* sp. P-121-IIcd ITS (AJ969550.1) as a query. We aligned the sequences using MAFFT v7.271 (Katoh & Standley, 2013) and trimmed the alignment to remove positions absent in over 20% of sequences using trimAl v1.2 (Capella-Gutiérrez *et al*., 2009). The Maximum-Likelihood phylogenetic tree was constructed with IQ-TREE v2.2.2.2 (Minh *et al*., 2020) using 10,000 rapid bootstraps. To construct a phylogenomic tree and carry out orthogroups analysis, we combined 23 reference genomes from literature (Table S2) and the predicted proteomes from our genome assembly and *Trebouxia* sp. SL0000003, and analyzed them with OrthoFinder v2.5.4 (Emms & Kelly, 2019). Phylogenies were visualized using iTOL v7 (Letunic & Bork, 2024).

### Horizontal gene transfer (HGT) analysis

To identify the HGT candidates reported by Beck *et al*. (2015), we used their nucleotide sequences (KF573967, KF573968, KF573969) as a BLAST query. The identified proteins were searched against NCBI non redundant protein (nr) and nucleotide (nt) databases (access date 2025/04/15). We ran three searches: (a) including all taxa; (b) excluding fungi; (c) including only Viridiplantae hits. We collected the top 100 hits from all searchers and excluded all hits with query coverage below 75%. The remaining protein sequences were combined with sequences from *T.* sp. SL0000003 and *T.* sp. ‘A48’ proteomes. We aligned the sequences and reconstructed phylogeny as described above.

### Microscopy

To visualize the *Trebouxia* strain, we used a Leica SP8 confocal microscope. Chlorophyll autofluorescence was excited at 514 nm and detected at 675-750 nm. Images were processed in ImageJ (Schneider *et al*., 2012).

## Results

### The genome sequence of *Trebouxia* sp. ‘A48’

Here we present a chromosome-level assembly of a *Trebouxia* alga (Figure 1a-b), which was isolated from the lichen thallus of *Xanthoria parietina* (Figure 1c). The nuclear genome consists of 69.1 Mbp was recovered in 20 contigs, of which 16 have telomeric repeats at both ends and likely represent complete chromosomes (Figure 2a, Table 1). The genome is 97.2% complete according to BUSCO and has a duplication rate of 0.6% (Figure 2b). Both plastid and mitochondrial genomes were recovered as single circular contigs, 320.5 and 109.5 kbp, respectively (Figure 2c-d). Repeats accounted for 12.72% of the genome and included chiefly LINE elements (2.75%), simple repeats (1.82%), LTR elements (1.42%), and unclassified repeats (5.64%). *De novo* annotation of the nuclear genome produced 12,820 gene models. The genome size, repeat content, and the number of gene models are consistent with recently published genome sequences of *Trebouxia* (Poquita-Du *et al*., 2024).

**Figure 2.**
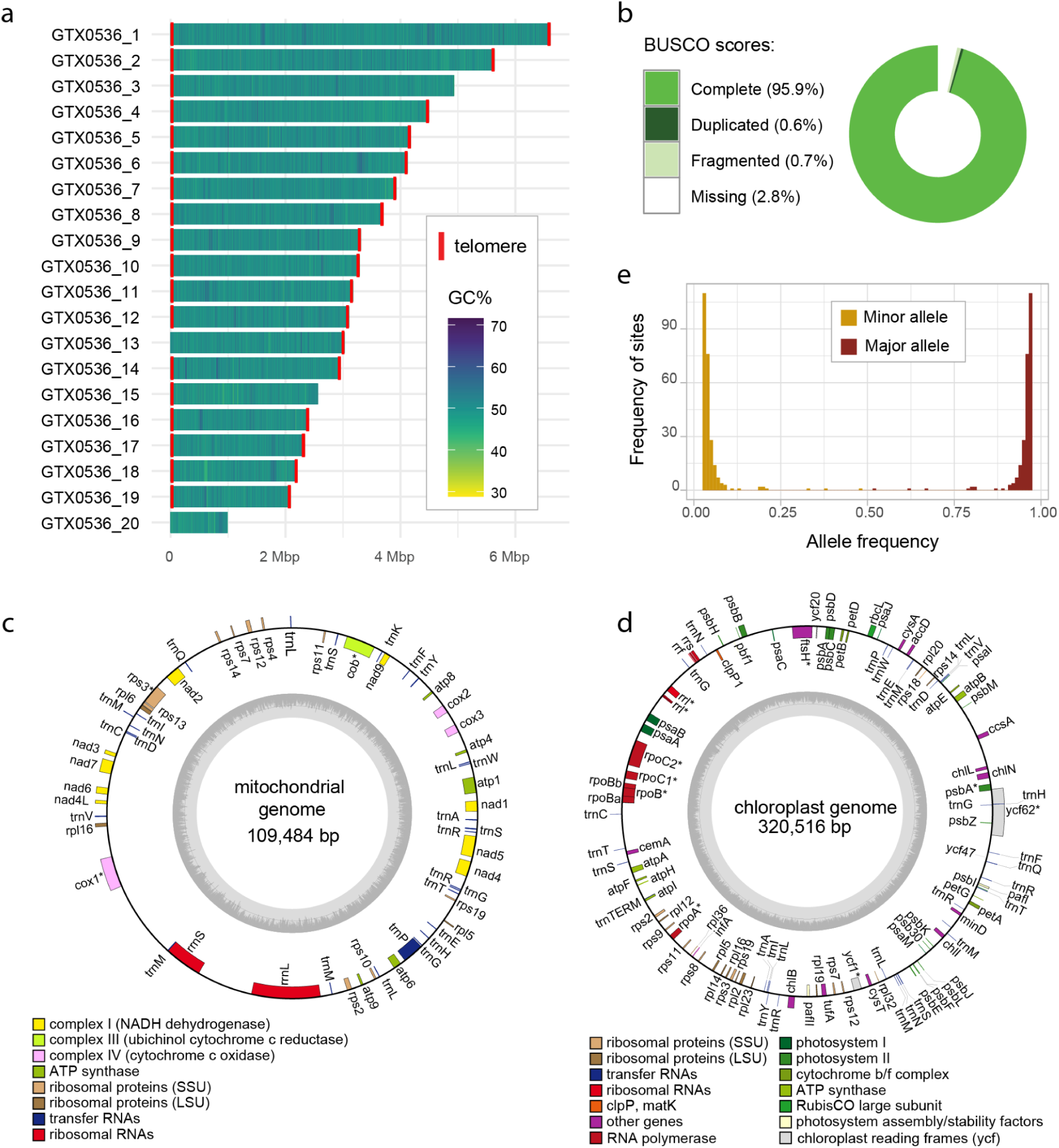
The genome of *Trebouxia* sp. ‘A48’. **A.** Plot showing the number, length, and GC-content of contigs comprising the nuclear genome. Red bars represent telomeric repeats. **B.** BUSCO quality scores (chlorophyta_odb10 database). **C.** Mitochondrial genome. The inner circle shows the GC-content. The genes are colored according to their function; the genes shown in the inner side of the circle are transcribed clockwise; the genes on the outside are transcribed counter-clockwise. **D.** Chloroplast genome. **E.** Allele frequency distribution in the nuclear genome. We removed positions within repeat elements and those outside of the coverage depth window 200-240x. The graph only shows allelic distributions between 0.03 and 0.97.

**Table 1.** Characteristics of the genomic assembly of *Trebouxia* sp. ‘A48’ compared to *Trebouxia* sp. SL0000003.

| Characteristic | *Trebouxia* sp. ‘A48’ | *Trebouxia* sp. SL0000003 |
| --- | --- | --- |
| Genome size, bp | 69,092,103 | 63,705,246 |
| Contig number in nuclear genome | 20 | 20 |
| Contigs with telomeric repeats on both ends | 16 | 3 |
| N50, Mbp | 3.71 | 1.19 |
| BUSCO completeness | 97.2% | 92.8% |
| BUSCO duplication | 0.6% | 0.7% |
| Repeat content | 12.72% | 10.6% |
| GC content | 49.68% | 50.3% |
| Number of genes | 12,820 | 10,061 |
| Number of proteins | 14,492 | 9,928 |
| Source | This study | Aquatic Symbiosis Genomic project |
| Source type | Isolate | Metagenome |
| Accession Number | Pending | PRJEB86095 |

The alga was identified as a not yet formally described species *T.* sp. ‘A48’ from the *T. decolorans* species complex, which belongs to clade ‘A’ within the *Trebouxia* genus (Muggia *et al*., 2020) (Figure 3a-b). Our isolate is closely related to photobionts of *Xanthoria* lichens collected from various geographic locations; its sister samples originated from Australia (AJ969551) and France (AJ969562) (Nyati *et al*., 2014).

**Figure 3.**
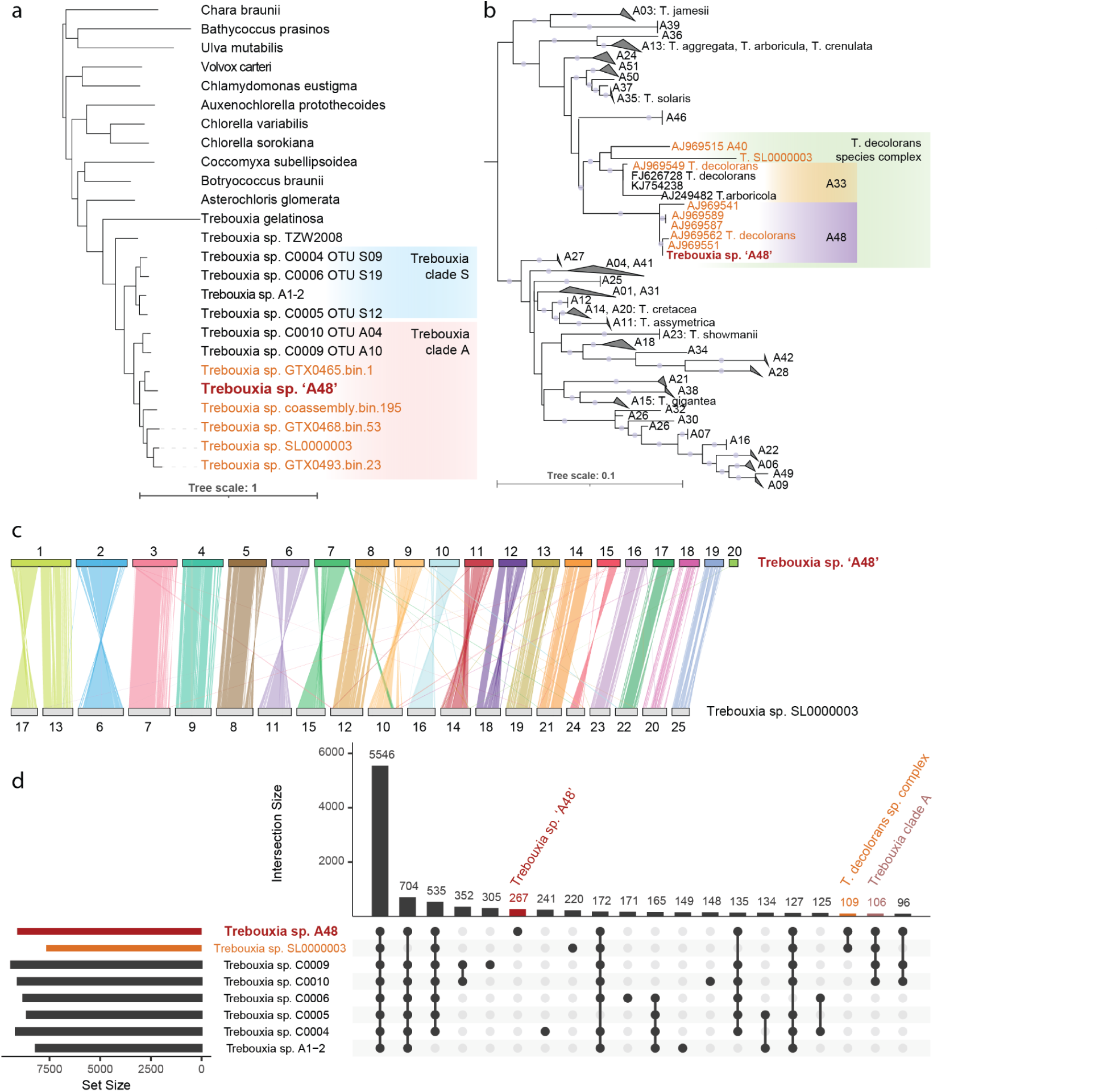
Evolutionary context of *Trebouxia* sp. ‘A48’. **A.** Species tree generated with Orthfinder based on 333 genomic loci. *Trebouxia* sp. ‘A48’ is shown in red; the metagenome-assembled genomes of *Trebouxia* from *X. parietina* lichens are shown in orange. The genomes from the literature are listed in Table S2; the tree in Newick format can be found in GitHub (https://github.com/metalichen/2025-Trebouxia-genome). **B.** ITS-based maximum-likelihood phylogeny of *Trebouxia*. Only a portion of the tree is shown; the entire tree in Newick format can be found in GitHub (https://github.com/metalichen/2025-Trebouxia-genome). The sequences used in the analysis are listed in Table S1. *Trebouxia* sp. ‘A48’ is shown in red; other *Trebouxia* from *Xanthoria* species are shown in orange (only shown for the *T. decolorans* clade). The clades are labeled with the OTU names as defined by Muggia *et al*. (2020). Dots represent bootstrap support >95%. **C.** Synteny between the genome of *Trebouxia* sp. ‘A48’ and the metagenome-assembled genome of *Trebouxia* sp. SL0000003 (PRJEB86095) isolated from a *X. parietina* metagenome. **D.** Upset plot showing intersection of orthogroups between *Trebouxia* species. We excluded from this graph metagenome-assembled genome from Tagirdzhanova *et al*. (2025), as they had lower completeness, and *T.* sp. TZW2008 and *T. gelatinosa*, for which we had only BUSCO annotations. Only top 20 sets are shown.

### *Trebouxia* sp. ‘A48’ in the context of its genus

We compared the *T.* sp ‘A48’ genome to the only other chromosome-level assembly of *Trebouxia*, which was generated as part of the Aquatic Symbiosis Genomics project (Table 1). This assembly was extracted from a long-read metagenome of *X. parietina* using binning and HiFi data (McKenna *et al*., 2024) and deposited in NCBI as *T.* sp. SL0000003. Our phylogenetic analysis recovered *T.* sp. SL0000003 as another member of the *T. decolorans* species complex and a sister to ‘A40’ and ‘A33’ OTUs (Figure 3b). The two assemblies exhibit significant synteny. Each contig from the *T.* sp. ‘A48’ assembly, for example, corresponded to one contig in *T.* sp. SL0000003, with only two exceptions (Figure 3c). First, the *T.* sp. ‘A48’ contig #1 was ‘split’ between two *T.* sp. SL0000003 contigs (OZ234913 and OZ234917), but whether this resulted from different genomic architectures or from fragmentation of the latter assembly is not clear. Second, the *T.* sp. ‘A48’ contig #20 did not have a match in the *T.* sp. SL0000003 genome, which might explain the lower completeness score received by *T.* sp. SL0000003. In addition, structural changes were present in contigs #5 and #12. The *Trebouxia* sp. ‘A48’ genome is larger, has a higher repeat content and more genes (Table 1).

The two genomes shared 7,222 orthogroups, with 1,864 families unique to *T.* sp. ‘A48’ and 448 unique to *T.* sp. SL0000003. However, the absence of some gene families from *T.* sp. SL0000003 might have resulted from the lower assembly completeness. By comparing *T.* sp. ‘A48’ to other *Trebouxia* species, we identified 267 gene families unique to this isolate (Figure 3d). In total, 109 orthogroups were present in both *T.* sp. ‘A48’ and *T.* sp. SL0000003 and no other genomes (Table S3). These included three orthogroups (with six *T.* sp. ‘A48’ genes) of various protease-encoding genes, one orthogroup containing three putative transporters with unknown functions, one predicted lectin, and one putative chitinase with a LysM domain.

### Genomic evidence for life cycle characteristics of *Trebouxia*

Allelic frequency distribution of the genome indicates that *T.* sp A48 is haploid (Figure 2e). While this result is consistent with the assumption frequently made regarding lichen algae (Fernández-Mendoza *et al*., 2011; Dal Grande *et al*., 2013; Greshake Tzovaras *et al*., 2020), it stands in contrast to recent reports that *T. lynnae*, another member of clade ‘A’, has a diplontic life cycle (Gazquez *et al*., 2024). Similarly to *T. lynnae,* the genome of *Trebouxia* sp. ‘A48’ encodes meiosis and mating-associated genes, including the nuclear fusion gene *GEX1*, gamete fusion gene *HAP2*, meiotic recombination protein *DMC1*, meiotic nuclear division gene *MND1*, and the meiosis specific protein *ZIP4*, but not *HOP1*, which was previously reported to be missing from another Trebouxiales genome (Fučíková *et al*., 2015) (Table S4, Table S5). In addition, *Trebouxia* sp. ‘A48’ genome encoded the meiosis-specific gene *HOP2*, which was lacking in *T. lynnae*. Across the genome, we identified 42 genes encoding flagellum-associated proteins (Table S5).

Some Volvocine algae, including *Chlamydomonas*, have well-documented heterothallism powered by sex-determining regions located on UV sex chromosomes (Yamamoto *et al*., 2021). However, nothing is known about possible mating types in trebouxiophycean algae. We therefore searched the *Trebouxia* sp. ‘A48’ genome for mating-type specific genes using sequences from *C. reinhardtii*, but failed to locate both MT+ specific *FUS1* and MT- specific *MTD1*. The genome encodes five RWP-RK transcription factor genes (Table S4), which are implicated in sex determination in diverse chlorophyte algae (Coelho *et al*., 2018), but their roles remain unclear.

### Genes involved in carbon-concentrating mechanisms in *Trebouxia*

Carbon-concentrating mechanisms (CCM) is a strategy for increasing photosynthesis efficiency common in eukaryotic algae and relatively rare in land plants (Maberly & Gontero, 2017). In chlorophyte algae, CCM relies on pyrenoids, which are organelles residing in chloroplasts. Not all lichen photobionts possess CCM, and their presence or absence can influence the geographic ranges of lichen species (Koch *et al*., 2023). Pyrenoids are present in *Trebouxia* algae, and their morphology has been long used as a diagnostic trait for distinguishing between different species (Bordenave *et al*., 2022). In the genome of *Trebouxia*, we found both key genes groups involved in pyrenoid-based CCM (He *et al*., 2023): bestrophin-like channels, which transport HCO_3_ ^−^ into the thylakoid lumen, and carbonic anhydrases, which catalyze conversion between CO_2_ and HCO_3_ ^−^ (Table S5). Three putative bestrophin-like channel genes (GTX0536PRED_011525-T1, GTX0536PRED_011526-T1, and GTX0536PRED_011527-T1) were located next to each other on contig GTX0536_17 separated by only 1,534 and 2,539 bp respectively. While two modes of CCM are identified in *Chlamydomonas reinhardtii*, we were able to find genes associated with only one. Namely, it was active chloroplast HCO_3_ ^−^ uptake, which in *C. reinhardtii* only occurs in very low CO_2_ conditions (He *et al*., 2023). In the *Trebouxia* genome, we found genes similar to transporters *HLA3* and *LCIA*, which move HCO_3_ ^−^ across the plasma membrane and the chloroplast envelope respectively (Table S5).

In addition, we found multiple genes involved in C4 metabolism, the CCM of land plants. These genes included phosphoenolpyruvate carboxylase *PEPC*, phosphoenolpyruvate carboxykinase *PCK*, and others (Figure S2), which is consistent with a previous report on *Trebouxia* (Alberola, 2015). However, these genes are widespread across many eukaryotes including non-photosynthetic lineages, and their presence alone cannot indicate C4 metabolism (Chi *et al*., 2014).

### Horizontally transferred genes in *Trebouxia*

We confirmed two instances of ancient horizontal gene transfer (HGT) from fungi to the *T.* sp. ‘A48’ genome. Previously, Beck *et al*. (2015) reported three putative HGT candidates from a *T. decolorans* genome. We were able to find two of them in the *T*. sp. A48 genome sequence: an oxidoreductase-like gene (GTX0536PRED_004195) and a sulfite efflux pump/TDT transporter-like gene (GTX0536PRED_006449). Since many more algal genomes are available now compared to the date of the original publication, we decided to proof-test the HGT reports. This yielded results largely consistent with Beck *et al*. (2015). When searched against the NCBI nr and nt databases, the sequences returned mostly fungal hits (Figure 4a, Table S6). For comparison, we analyzed two genes immediately upstream of the HGT candidates, which returned no fungal hits at all (Figure 4b). In the phylogenies, both candidate HGT genes were recovered in chlorophyte clades nested within the larger fungal clade (Figure 4c-d). In both cases, the sequences were recovered not in the lecanoromycete clade, as would be expected for a mycobiont-to-photobiont HGT, but in the basal part of the fungal clade, which indicates a more ancient transfer. The genome also encodes two enzymes from the CAZy family GH8 (Table S4), which were recently revealed as potentially horizontally transferred to Trebouxiophyceae from bacteria (Puginier *et al*., 2024).

**Figure 4.**
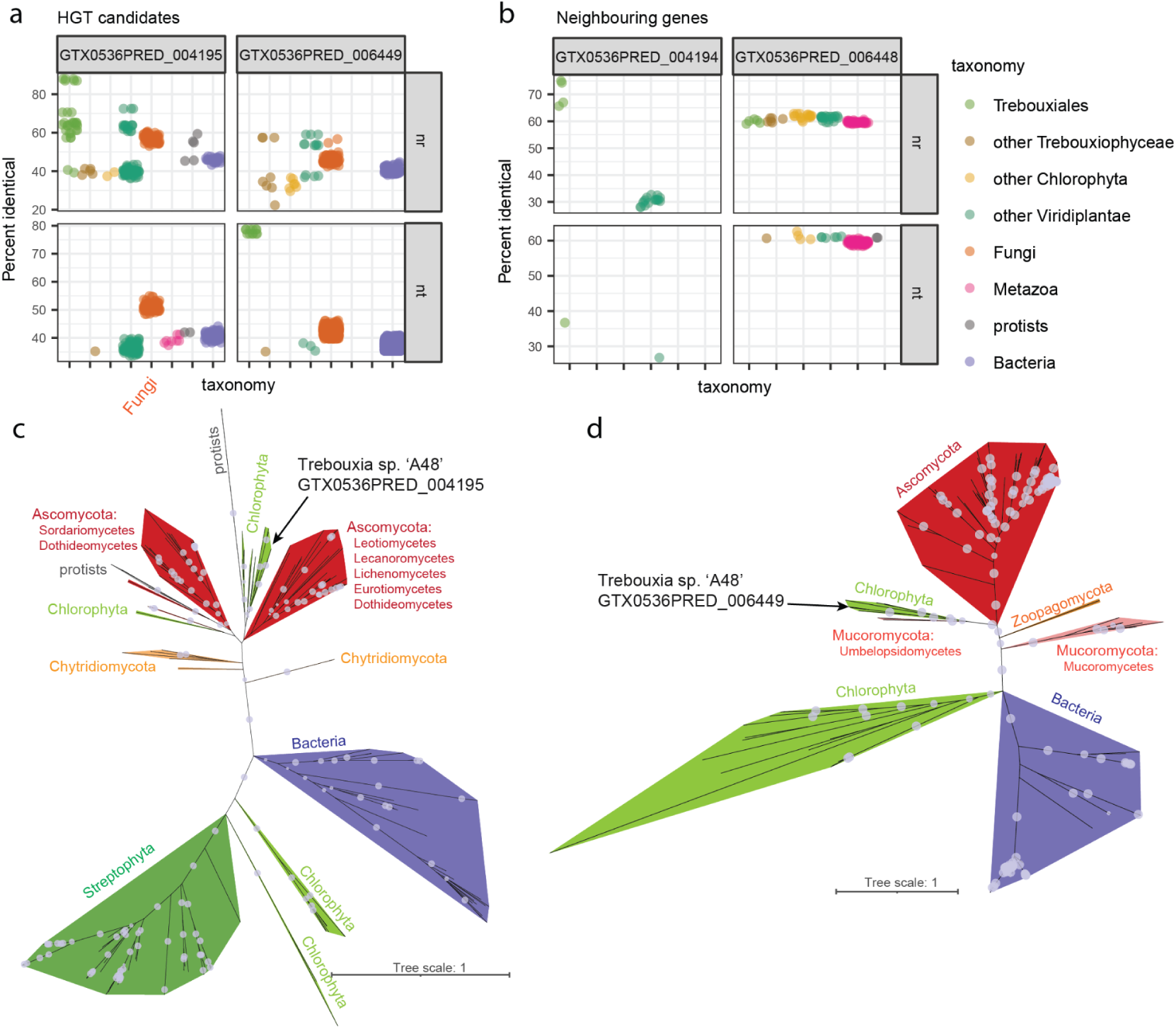
Predicted instances of horizontal gene transfer (HGT) in the *Trebouxia* sp. ‘A48’ genome. A-B. Results of a BLAST search to the NCBI nt and nr databases for the HGT candidate (a) and their neighboring genes (b). Each dot represents a hit, positioned based on the taxonomy of the subject sequence and the percent identity of the hit. Only hits with the query coverage above 75% are shown. **C-D.** Phylogenetic trees derived from a Maximum-Likelihood analysis of homologues for the HGT candidates GTX0536PRED_004195 (c) and GTX0536PRED_006449 (d). Dots represent bootstrap support >90%. The trees in Newick format can be found in GitHub (https://github.com/metalichen/2025-Trebouxia-genome). The sequences used in the analysis are listed in Table S6.

### Secretome of *T.* sp. ‘A48’ is enriched in hydrolases and redox enzymes

Algal secreted proteins have been implicated in both symbiosis and the desiccation response of Trebouxiophyceae algae (Armaleo *et al*., 2019; González-Hourcade *et al*., 2021). Using three bioinformatics tools (SignalP, deep TMHMM, and WolfPSORT), we identified 103 proteins as being putatively secreted by *Trebouxia* (Table S7). The secretome size is consistent with those reported from other Trebouxiophyceae (González-Hourcade *et al*., 2021). More than half of the predicted secreted proteins lacked any functional annotation (Figure 5a), but among the 46 that did, several functional groups were overrepresented, including hydrolases and redox proteins (Figure 5b).

**Figure 5.**
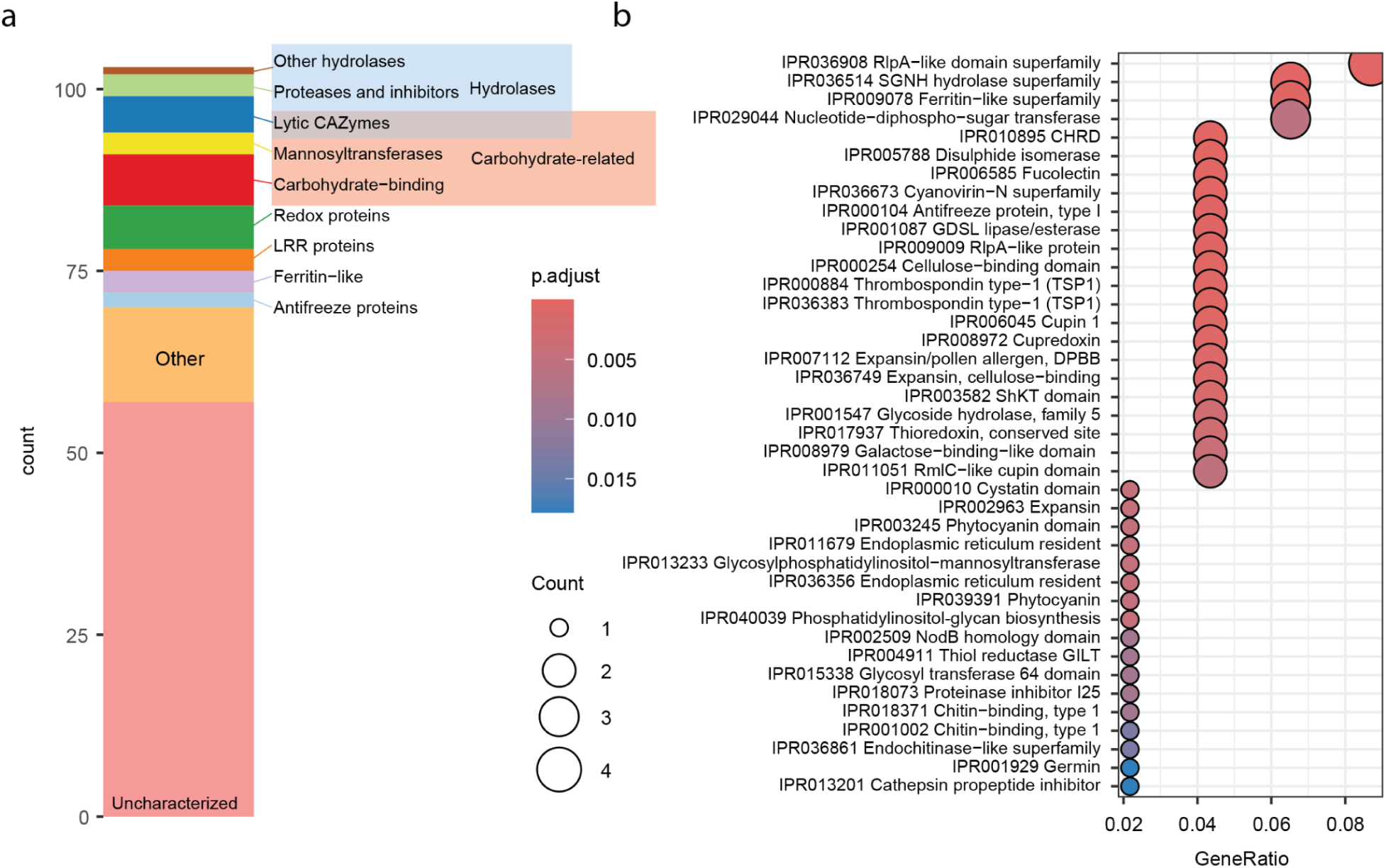
The predicted secretome of *Trebouxia* sp. ‘A48’. **A.** Composition of the secretome. The 103 secreted genes were predicted *in silico* and grouped based on their functional annotation. LRR stands for leucine-rich repeat; CAZyme stands for carbohydrate-active enzyme. **B.** Enrichment analysis showing InterProScan domains overrepresented in the secretome. The size of the dots represents the number of proteins; the color represents p-value.

Fungal lectins have long been hypothesized to play a role in lichen symbiont recognition and adhesion, but less is known regarding photobiont-produced lectins (Singh & Walia, 2014). We found that carbohydrate-related proteins, including enzymes, accounted for 17% of the predicted secretome. The secretome contained one endochitinase-like glycoside hydrolase, for example, which might target fungal cell walls. Seven secreted proteins were also predicted to bind carbohydrates; those included three expansin-like proteins, two of which also contained an RlpA-like domain, a Ricin B lectin-like and a fucolectin-like proteins. Three alpha-mannosyltransferases from CAZy families GT32, GT64, and GT71 were detected too, which is notable given the abundance of alpha-mannans in lichen thalli (Spribille *et al*., 2020).

As oxidative stress accompanies desiccation, lichen symbionts possess mechanisms for protection against reactive oxygen species (Kranner *et al*., 2003). Among the redox proteins from lichen algae, one predicted secreted protein has previously been implicated in the desiccation response (González-Hourcade *et al*., 2021). In the secretome, we identified several proteins similar to cupredoxin, thioredoxin, and manganese/iron superoxide dismutase (Table S7).

Notably, the secretome also contained two proteins annotated as antifreeze protein 1-like (Figure 5). This functional family was originally described from marine animals, and while similar proteins appear in chlorophyte algae (e.g. Cre09.g398150 in *C. reinhardtii*), their role remains unknown.

Among the functionally characterized secreted proteins, many shared annotations with gene families identified by Puginier *et al*. (2024) as associated with lichenization in chlorophyte algae. These included ferritin, fucolectin, and cupin-like proteins. In general, the contents of the predicted secretome appeared rather conserved when compared to other *Trebouxia* species.

Nearly half of the predicted secreted proteins (n=46), for example, were encoded by genes from orthogroups shared between all eight surveyed genomes. The conserved portion of the secretome included 59% of all functionally characterized proteins and all but one predicted hydrolase. By contrast, nine proteins were encoded by genes from orthogroups unique to *T.* sp. ‘A48’, only one of them had a leucine-rich repeat domain and the rest lacked functional annotation. One additional secreted protein belonged to an orthogroup unique for *T. decolorans* species complex; it too had no functional annotation.

## Discussion

Lichen symbioses constitute a wide array of associations between vastly different fungi and photosynthetic organisms united by a similar outcome in the complex form of a multicellular lichen thallus (Spribille *et al*., 2022). Remarkably, the lichen thallus shows considerable anatomical complexity that is strongly heritable, but results from the association of morphologically simple fungal, cyanobacterial or algal partners. The complexity of partners in lichens is also increasingly apparent with consistent presence of bacteria and non-mycobiont fungi reported from several lichens, including *Xanthoria parietina* (Hodkinson & Lutzoni, 2009; Grimm *et al*., 2021; Leiva *et al*., 2021; Tagirdzhanova *et al*., 2024, 2025). However, the majority of biomass in a lichen is typically represented by a major mycobiont fungus and a photosynthetic partner. In this regard, *Trebouxia* algae play an outsized role in lichen biology as a whole, because they participate in the most species-rich groups of lichens, formed by mycobionts from orders Lecanorales including Parmeliaceae, Caliciales, and Teloschistes, and are also paired with a diversity of mycobionts from classes Lecanoromycetes, Lichinomycetes, and Arthoniomycetes (Sanders & Masumoto, 2021). Investigating the biology of *Trebouxia* is therefore pivotal to understanding lichen physiology and development.

In this report, we generated a chromosome-level assembly of the genome of *T.* sp ‘A48’ from the *T. decolorans* species complex. This group of species is primarily associated with the mycobionts of the Teloschistales order, but is also reported from lichens formed by Lecanorales and Caliciales fungi (Dal Grande *et al*., 2014; Werth & Sork, 2014). We used the genome to investigate the evolutionary history of the alga and its relationship with its fungal partner. We have confirmed two instances of ancient horizontal gene transfer from a fungal origin and also profiled the predicted secretome of the alga. We identified genes that likely play a role in carbon concentration – an important physiological mechanism that allows algae to withstand low CO_2_ conditions. CCM appears to be especially important for lichens exposed to direct light and liquid water (Koch *et al*., 2022), which is often the case for *X. parietina*. The presence of CCM in *Trebouxia* might be one of the reasons for its prevalence in lichen symbioses.

Fungal secreted proteins play a crucial role in both mutualistic and parasitic relationships (Lowe & Howlett, 2012) and lichen fungi are likely no exception (Snelders *et al*., 2022; Tagirdzhanova *et al*., 2025). Predicted effector proteins are, for example, found in *X. parietina* that are potentially deployed by the mycobiont during lichen development, perhaps to manipulate its photobiont partner. Much less is known, however, about the secretomes of chlorophyte algae, but existing studies suggest that algae secrete proteins though both conventional and unconventional exocytotic pathways, and that their secretomes are dominated by extracellular enzymes (Xiao & Zheng, 2016; Choi *et al*., 2021). A recent study used proteomics and genomics to profile secretomes of two lichen algae, including *Trebouxia lynnea* (González-Hourcade *et al*., 2021). *T.* sp ‘A48’ has a similar secretome profile to *T. lynnea*, and both secretomes contain lectins and CAZymes potentially involved in symbiotic interactions or cell adhesion.

Our results also provide evidence that *T.* sp ‘A48’ is haploid and capable of sexual reproduction, as its genome encodes all key genes required for meiosis and mating. At the same time, it is unclear to what extent this potential is realized in nature. Marker-gene based analysis of the genetic structure of *X. parietina* photobionts showed low admixture rates, consistent with *T. decolorans* existing primarily as a clonal organism (Wyczanska *et al*., 2023). Notably, while examining the *T.* sp. ‘A48’ culture, we detected flagellated cells highly similar to those identified as gametes in *T. lynnae* (Gazquez *et al*., 2024) (Figure 1b), which is consistent with the presence of flagellum-associated genes in the *T.* sp. ‘A48’ genome. However, these cells might represent asexual zoospores.

Arguably, the main obstacle to understanding the lichen symbiosis is our inability to re-create the symbiosis from its constituent parts under controlled conditions. Lichen studies instead most often rely on natural lichen thalli collected from the field, which inevitably introduces a wide array of variables beyond the researchers’ control. One such factor is the identity of the symbionts, which in lichens often resemble a rotating cast of players rather than a straightforward one-to-one relationship (Spribille *et al*., 2022). Lichen mycobionts tend to associate with several closely related photobiont lineages, while the photobionts tend to be less restrictive (Sanders & Masumoto, 2021). The mechanisms for lichen symbiont recognition, however, remain largely unknown. In this study we compared the genomes of two *Trebouxia* photobionts of *X. parietina* lichens. Despite belonging to different candidate species, the two genomes were similar in both genomic structure and gene content. We attempted to identify the key features of the two genomes that distinguish them from other *Trebouxia* species that are not known from *Xanthoria* lichens. Among the genes present in both genomes but not found in any other studied *Trebouxia*, two candidates encoded proteins that can selectively bind polysaccharides, hinting at a potential role in symbiont recognition. Neither of these genes were identified as secreted, however, although it is possible that they are secreted through unconventional routes. Given the fact that the *X. parietina* symbiosis is established *de novo* every generation from its constituent fungal and algal partners, the photobiont composition is remarkably consistent across the wide geographic area inhabited by this lichen. The ITS sequence of our UK-originated strain was, for instance, nearly identical to those of *X. parietina* photobionts from France and Australia, and highly similar to those from North America. Given this degree of conservation, the genome sequence generated here may provide a valuable reference for studies on *X. parietina* regardless of the geographic location of collected lichens.

In summary, we set out to produce a reference genome for the photobiont of *X. parietina*, which can be considered a model lichen symbioses. With an N50 of 3.7 Mbp and 16 contigs out of 20 likely representing telomere-to-telomere chromosome assemblies, this genome is, to our knowledge, the most contiguous and complete genome of a lichen alga currently available. The corresponding algal strain is available to the community and can be obtained via the SAG algal culture collection. We hope that together these resources will aid research into chlorophyte algae genomics and evolution, as well as experimental studies of the lichen symbiosis.

## Acknowledgments

This work was supported by grants from The Halpin Family, The Gatsby Charitable Foundation, and the Biotechnology and Biological Sciences Research Council BBS/E/J/000PR9798 to NJT. The authors thank Maike Lorenz (SAG) for help with depositing the algal strain and Alison MacFadyen for help with depositing data.

## Author contributions

GT and NJT planned and designed the study. JR and GT performed lab work. GT performed bioinformatic analysis and microscopy and drafted the manuscript. All authors contributed to editing.

## Data and code availability

The genome assembly and annotation and the raw data will be made publicly available (submission pending the assignment of a permanent SAG collection number). All scripts created for this project and detailed information on the usage of bioinformatics software are available at GitHub (https://github.com/metalichen/2025-Trebouxia-genome).

## Supplementary images

**Figure S1.**
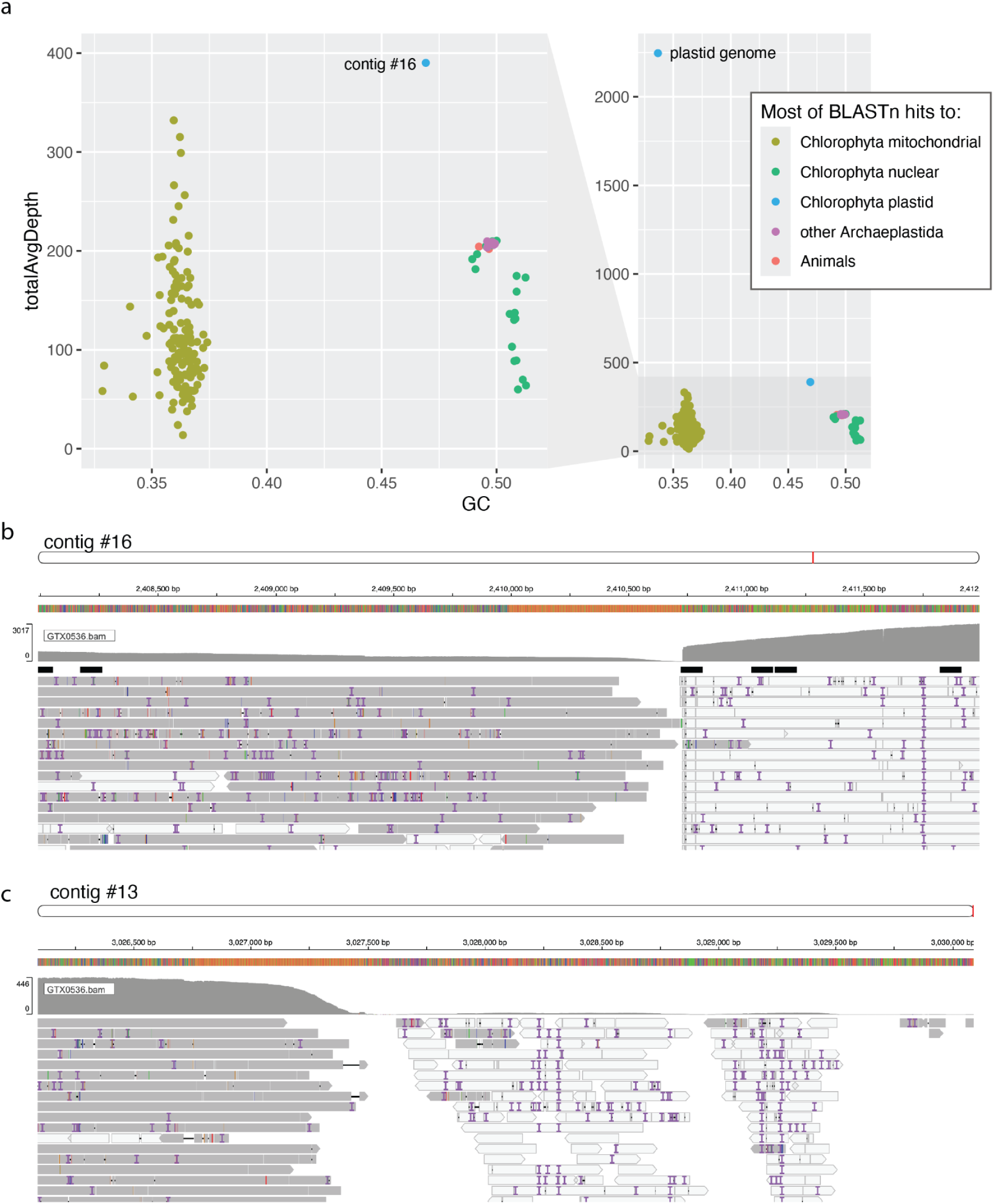
Assembly of *Trebouxia* sp. “A48” genome. **A.** GC/coverage plot of the entire assembly. Each dot represents a contig colored based on its majority hits to the NCBI nt database. The y-axis corresponds to the coverage depth averaged across the contig, the x-axis corresponds to the GC content. The plastid genome was recovered as a single circular contig. The contigs with mitochondrial hits accounted for the majority of contigs in the assembly; these data were subsequently re-assembled to obtain a single contig assembly of the mitochondrial genome. **B.** Misassembly in the contig #16 visualized with IGV. The top track shows the position of the highlighted region within the contig. The track below shows the sequence of the region. The histogram shows the coverage depth for each position. The bottom track shows a portion of reads aligned to this region. Contig #16 was previously shown to have coverage than the rest of the nuclear genome and majority of hits to plastid sequences (Figure S1a) This anomaly resulted from a misassembly, in which a nuclear contig was fused with a higher-coverage plastid contig. **C.** Misassembly in the terminal region of contig #13 visualized as described above.

**Figure S2.**
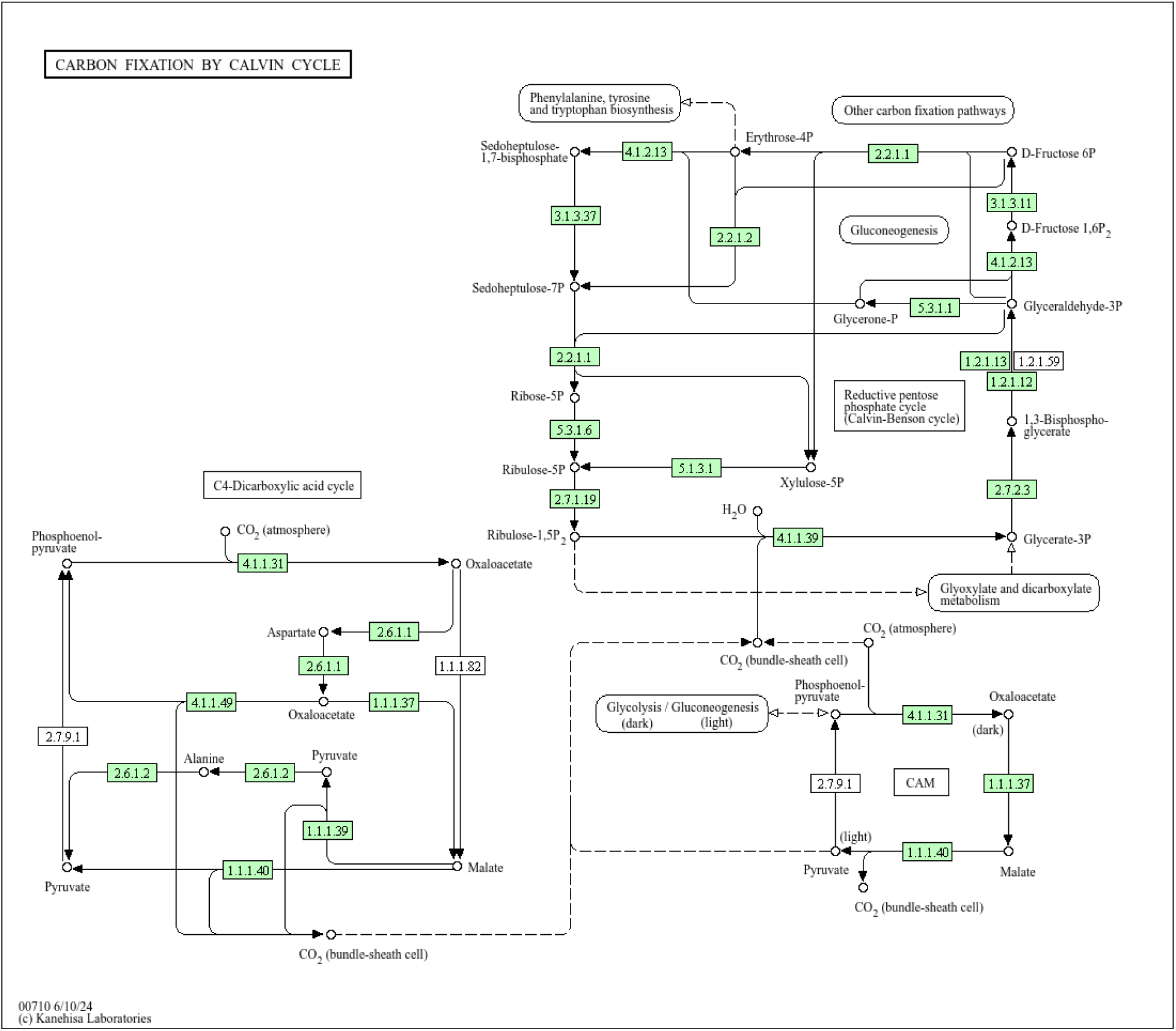
Metabolic map showing genes involved in carbon fixation and C4 metabolism. Green blocks correspond to genes identified in the genome; white blocks represent absent genes. The map is generated using the KEGG Mapper Reconstruct tool.

## References

Alberola FM. 2015. Caracterización genómica del microalga Trebouxia sp.TR9 aislada de liquen Ramalina farinacea (L.) Ach. mediante secuenciación masiva.

Almagro Armenteros JJ, Tsirigos KD, Sønderby CK, Petersen TN, Winther O, Brunak S, von Heijne G, Nielsen H. 2019. SignalP 5.0 improves signal peptide predictions using deep neural networks. Nature biotechnology 37: 420–423.

Ament-Velásquez SL, Tuovinen V, Bergström L, Spribille T, Vanderpool D, Nascimbene J, Yamamoto Y, Thor G, Johannesson H. 2021. The Plot Thickens: Haploid and Triploid-Like Thalli, Hybridization, and Biased Mating Type Ratios in. Frontiers in fungal biology 2: 656386.

Armaleo D, Müller O, Lutzoni F, Andrésson ÓS, Blanc G, Bode HB, Collart FR, Dal Grande F, Dietrich F, Grigoriev IV, et al. 2019. The lichen symbiosis re-viewed through the genomes of Cladonia grayi and its algal partner Asterochloris glomerata. BMC genomics 20: 605.

Beck A, Divakar PK, Zhang N, Molina MC, Struwe L. 2015. Evidence of ancient horizontal gene transfer between fungi and the terrestrial alga Trebouxia. Organisms, diversity & evolution 15: 235–248.

Bordenave CD, Muggia L, Chiva S, Leavitt SD, Carrasco P, Barreno E. 2022. Chloroplast morphology and pyrenoid ultrastructural analyses reappraise the diversity of the lichen phycobiont genus Trebouxia (Chlorophyta). Algal research 61: 102561.

Camacho C, Coulouris G, Avagyan V, Ma N, Papadopoulos J, Bealer K, Madden TL. 2009. BLAST+: architecture and applications. BMC bioinformatics 10: 421.

Capella-Gutiérrez S, Silla-Martínez JM, Gabaldón T. 2009. trimAl: a tool for automated alignment trimming in large-scale phylogenetic analyses. *Bioinformatics (Oxford*, England) 25: 1972–1973.

Chan PP, Lowe TM. 2019. tRNAscan-SE: Searching for tRNA Genes in Genomic Sequences. Methods in molecular biology 1962: 1–14.

Cheng H, Concepcion GT, Feng X, Zhang H, Li H. 2021. Haplotype-resolved de novo assembly using phased assembly graphs with hifiasm. Nature methods 18: 170–175.

Chi S, Wu S, Yu J, Wang X, Tang X, Liu T. 2014. Phylogeny of C4-photosynthesis enzymes based on algal transcriptomic and genomic data supports an archaeal/proteobacterial origin and multiple duplication for most C4-related genes. PloS one 9: e110154.

Choi J, Shin J-H, An HJ, Oh MJ, Kim S-R. 2021. Analysis of secretome and N-glycosylation of Chlorella species. Algal research 59: 102466.

Coelho SM, Gueno J, Lipinska AP, Cock JM, Umen JG. 2018. UV Chromosomes and Haploid Sexual Systems. Trends in plant science 23: 794–807.

Dal Grande F, Alors D, Divakar PK, Bálint M, Crespo A, Schmitt I. 2014. Insights into intrathalline genetic diversity of the cosmopolitan lichen symbiotic green alga Trebouxia decolorans Ahmadjian using microsatellite markers. Molecular phylogenetics and evolution 72: 54–60.

Dal Grande F, Beck A, Singh G, Schmitt I. 2013. Microsatellite primers in the lichen symbiotic alga Trebouxia decolorans (Trebouxiophyceae). Applications in plant sciences 1.

Dobin A, Davis CA, Schlesinger F, Drenkow J, Zaleski C, Jha S, Batut P, Chaisson M, Gingeras TR. 2013. STAR: ultrafast universal RNA-seq aligner. *Bioinformatics (Oxford*, England) 29: 15–21.

Emms DM, Kelly S. 2019. OrthoFinder: phylogenetic orthology inference for comparative genomics. Genome biology 20: 238.

Fernández-Mendoza F, Domaschke S, García MA, Jordan P, Martín MP, Printzen C. 2011. Population structure of mycobionts and photobionts of the widespread lichen Cetraria aculeata. Molecular ecology 20: 1208–1232.

Fučíková K, Pažoutová M, Rindi F. 2015. Meiotic genes and sexual reproduction in the green algal class Trebouxiophyceae (Chlorophyta). Journal of phycology 51: 419–430.

Gazquez A, Bordenave CD, Montero-Pau J, Pérez-Rodrigo M, Marco F, Martínez-Alberola F, Muggia L, Barreno E, Carrasco P. 2024. From spores to gametes: A sexual life cycle in a symbiotic Trebouxia microalga. Algal research 84: 103744.

González-Hourcade M, Del Campo EM, Casano LM. 2021. The Under-explored Extracellular Proteome of Aero-Terrestrial Microalgae Provides Clues on Different Mechanisms of Desiccation Tolerance in Non-Model Organisms. Microbial ecology 81: 437–453.

Greiner S, Lehwark P, Bock R. 2019. OrganellarGenomeDRAW (OGDRAW) version 1.3.1: expanded toolkit for the graphical visualization of organellar genomes. Nucleic acids research 47: W59–W64.

Greshake Tzovaras B, Segers FHID, Bicker A, Dal Grande F, Otte J, Anvar SY, Hankeln T, Schmitt I, Ebersberger I. 2020. What Is in Umbilicaria pustulata? A Metagenomic Approach to Reconstruct the Holo-Genome of a Lichen. Genome biology and evolution 12: 309–324.

Grimm M, Grube M, Schiefelbein U, Zühlke D, Bernhardt J, Riedel K. 2021. The Lichens’ Microbiota, Still a Mystery? Frontiers in microbiology 12: 623839.

Haas BJ, Salzberg SL, Zhu W, Pertea M, Allen JE, Orvis J, White O, Buell CR, Wortman JR. 2008. Automated eukaryotic gene structure annotation using EVidenceModeler and the Program to Assemble Spliced Alignments. Genome biology 9: R7.

Hallgren J, Tsirigos KD, Pedersen MD, Almagro Armenteros JJ, Marcatili P, Nielsen H, Krogh A, Winther O. 2022. DeepTMHMM predicts alpha and beta transmembrane proteins using deep neural networks. bioRxiv.

He S, Crans VL, Jonikas MC. 2023. The pyrenoid: the eukaryotic CO2-concentrating organelle. The Plant cell 35: 3236–3259.

Hiltunen M, Ament-Velásquez SL, Johannesson H. 2021. The Assembled and Annotated Genome of the Fairy-Ring Fungus Marasmius oreades. Genome biology and evolution 13.

Hodkinson BP, Lutzoni F. 2009. A microbiotic survey of lichen-associated bacteria reveals a new lineage from the Rhizobiales. *Symbiosis (Philadelphia*, Pa.) 49: 163–180.

Honegger R. 1996. Experimental studies of growth and regenerative capacity in the foliose lichen *Xanthoria parietina*. The new phytologist 133: 573–581.

Honegger R, Zippler U, Scherrer S, Dyer PS. 2004. Genetic diversity in Xanthoria parietina (L.) Th. Fr. (lichen-forming ascomycete) from worldwide locations. *Lichenologist (London*, England) 36: 381–390.

Horton P, Park K-J, Obayashi T, Fujita N, Harada H, Adams-Collier CJ, Nakai K. 2007. WoLF PSORT: protein localization predictor. Nucleic acids research 35: W585–7.

Hua-Van A, Capy P. 2024. Transposable Elements and Genome Evolution. John Wiley & Sons.

Kang DD, Li F, Kirton E, Thomas A, Egan R, An H, Wang Z. 2019. MetaBAT 2: an adaptive binning algorithm for robust and efficient genome reconstruction from metagenome assemblies. PeerJ 7: e7359.

Katoh K, Standley DM. 2013. MAFFT multiple sequence alignment software version 7: improvements in performance and usability. Molecular biology and evolution 30: 772–780.

Knaus BJ, Grünwald NJ. 2017. vcfr: a package to manipulate and visualize variant call format data in R. Molecular ecology resources 17: 44–53.

Koboldt DC, Zhang Q, Larson DE, Shen D, McLellan MD, Lin L, Miller CA, Mardis ER, Ding L, Wilson RK. 2012. VarScan 2: somatic mutation and copy number alteration discovery in cancer by exome sequencing. Genome research 22: 568–576.

Koch NM, Lendemer JC, Manzitto-Tripp EA, McCain C, Stanton DE. 2023. Carbon-concentrating mechanisms are a key trait in lichen ecology and distribution. Ecology 104: e4011.

Koch NM, Stanton D, Müller SC, Duarte L, Spielmann AA, Lücking R. 2022. Nuanced qualitative trait approaches reveal environmental filtering and phylogenetic constraints on lichen communities. *Ecosphere (Washington*, D.C) 13.

Korf I. 2004. Gene finding in novel genomes. BMC bioinformatics 5: 59.

Korhonen P, Kallio P. 1987. The effect of different night conditions on the CO2 fixation in a lichen Xanthoria parietina. Photosynthesis research 12: 3–11.

Kranner I, Zorn M, Turk B, Wornik S, Beckett RP, Batič F. 2003. Biochemical traits of lichens differing in relative desiccation tolerance. The New phytologist 160: 167–176.

Lang BF, Beck N, Prince S, Sarrasin M, Rioux P, Burger G. 2023. Mitochondrial genome annotation with MFannot: a critical analysis of gene identification and gene model prediction. Frontiers in plant science 14: 1222186.

Leiva D, Fernández-Mendoza F, Acevedo J, Carú M, Grube M, Orlando J. 2021. The Bacterial Community of the Foliose Macro-lichen Peltigera frigida Is More than a Mere Extension of the Microbiota of the Subjacent Substrate. Microbial ecology 81: 965–976.

Letunic I, Bork P. 2024. Interactive Tree of Life (iTOL) v6: recent updates to the phylogenetic tree display and annotation tool. Nucleic acids research 52: W78–W82.

Li H. 2018. Minimap2: pairwise alignment for nucleotide sequences. *Bioinformatics (Oxford*, England) 34: 3094–3100.

Li H, Handsaker B, Wysoker A, Fennell T, Ruan J, Homer N, Marth G, Abecasis G, Durbin R, 1000 Genome Project Data Processing Subgroup. 2009. The Sequence Alignment/Map format and SAMtools. Bioinformatics (Oxford, England) 25: 2078–2079.

Lomsadze A, Ter-Hovhannisyan V, Chernoff YO, Borodovsky M. 2005. Gene identification in novel eukaryotic genomes by self-training algorithm. Nucleic acids research 33: 6494–6506.

Lowe RGT, Howlett BJ. 2012. Indifferent, affectionate, or deceitful: lifestyles and secretomes of fungi. PLoS pathogens 8: e1002515.

Maberly SC, Gontero B. 2017. Ecological imperatives for aquatic CO2-concentrating mechanisms. Journal of experimental botany 68: 3797–3814.

Majoros WH, Pertea M, Salzberg SL. 2004. TigrScan and GlimmerHMM: two open source ab initio eukaryotic gene-finders. Bioinformatics 20: 2878–2879.

McKenna V, Archibald JM, Beinart R, Dawson MN, Hentschel U, Keeling PJ, Lopez JV, Martín-Durán JM, Petersen JM, Sigwart JD, et al. 2024. The Aquatic Symbiosis Genomics Project: probing the evolution of symbiosis across the Tree of Life. Wellcome open research 6: 254.

Minh BQ, Schmidt HA, Chernomor O, Schrempf D, Woodhams MD, von Haeseler A, Lanfear R. 2020. IQ-TREE 2: New Models and Efficient Methods for Phylogenetic Inference in the Genomic Era. Molecular biology and evolution 37: 1530–1534.

Mistry J, Chuguransky S, Williams L, Qureshi M, Salazar GA, Sonnhammer ELL, Tosatto SCE, Paladin L, Raj S, Richardson LJ, et al. 2021. Pfam: The protein families database in 2021. Nucleic acids research 49: D412–D419.

Moriya Y, Itoh M, Okuda S, Yoshizawa AC, Kanehisa M. 2007. KAAS: an automatic genome annotation and pathway reconstruction server. Nucleic acids research 35: W182–5.

Muggia L, Nelsen MP, Kirika PM, Barreno E, Beck A, Lindgren H, Lumbsch HT, Leavitt SD, Trebouxia working group. 2020. Formally described species woefully underrepresent phylogenetic diversity in the common lichen photobiont genus Trebouxia (Trebouxiophyceae, Chlorophyta): An impetus for developing an integrated taxonomy. Molecular phylogenetics and evolution 149: 106821.

Nyati S, Scherrer S, Werth S, Honegger R. 2014. Green-algal photobiont diversity (Trebouxia spp.) in representatives of Teloschistaceae (Lecanoromycetes, lichen-forming ascomycetes). *Lichenologist (London*, England) 46: 189–212.

Palmer JM, Stajich J. 2020. Funannotate v1.8.1: Eukaryotic genome annotation. Zenodo.

Paysan-Lafosse T, Blum M, Chuguransky S, Grego T, Pinto BL, Salazar GA, Bileschi ML, Bork P, Bridge A, Colwell L, et al. 2023. InterPro in 2022. Nucleic acids research 51: D418–D427.

Poquita-Du RC, Otte J, Calchera A, Schmitt I. 2024. Genome-Wide Comparisons Reveal Extensive Divergence Within the Lichen Photobiont Genus, Trebouxia. Genome biology and evolution 16.

Puginier C, Libourel C, Otte J, Skaloud P, Haon M, Grisel S, Petersen M, Berrin J-G, Delaux P-M, Dal Grande F, et al. 2024. Phylogenomics reveals the evolutionary origins of lichenization in chlorophyte algae. Nature communications 15: 4452.

Quigley S, Damas J, Larkin DM, Farré M. 2023. syntenyPlotteR: a user-friendly R package to visualize genome synteny, ideal for both experienced and novice bioinformaticians. Bioinformatics advances 3: vbad161.

Rawlings ND, Waller M, Barrett AJ, Bateman A. 2014. MEROPS: the database of proteolytic enzymes, their substrates and inhibitors. Nucleic acids research 42: D503–9.

Sanders WB, Masumoto H. 2021. Lichen algae: the photosynthetic partners in lichen symbioses. *Lichenologist (London*, England) 53: 347–393.

Scharnagl K, Tagirdzhanova G, Talbot NJ. 2023. The coming golden age for lichen biology. Current biology : CB 33: R512–R518.

Schneider CA, Rasband WS, Eliceiri KW. 2012. NIH Image to ImageJ: 25 years of image analysis. Nature methods 9: 671–675.

Seppey M, Manni M, Zdobnov EM. 2019. BUSCO: Assessing Genome Assembly and Annotation Completeness. *Methods in molecular biology (Clifton*, N.J.) 1962: 227–245.

Singh RS, Walia AK. 2014. Characteristics of lichen lectins and their role in symbiosis. *Symbiosis (Philadelphia*, Pa.) 62: 123–134.

Snelders NC, Rovenich H, Thomma BPHJ. 2022. Microbiota manipulation through the secretion of effector proteins is fundamental to the wealth of lifestyles in the fungal kingdom. FEMS microbiology reviews 46.

Spribille T, Resl P, Stanton DE, Tagirdzhanova G. 2022. Evolutionary biology of lichen symbioses. The New phytologist 234: 1566–1582.

Spribille T, Tagirdzhanova G, Goyette S, Tuovinen V, Case R, Zandberg WF. 2020. 3D biofilms: in search of the polysaccharides holding together lichen symbioses. FEMS microbiology letters 367.

Stanke M, Waack S. 2003. Gene prediction with a hidden Markov model and a new intron submodel. Bioinformatics 19 Suppl 2: ii215–25.

Tagirdzhanova G, Saary P, Cameron ES, Allen CCG, Garber AI, Escandón DD, Cook AT, Goyette S, Nogerius VT, Passo A, et al. 2024. Microbial occurrence and symbiont detection in a global sample of lichen metagenomes. PLoS biology 22: e3002862.

Tagirdzhanova G, Scharnagl K, Sahu N, Yan X, Bucknell A, Bentham AR, Jégousse C, Ament-Velásquez SL, Onuț-Brännström I, Johannesson H, et al. 2025. Complexity of the lichen symbiosis revealed by metagenome and transcriptome analysis of Xanthoria parietina. Current biology : CB 35: 799–817.e5.

Tillich M, Lehwark P, Pellizzer T, Ulbricht-Jones ES, Fischer A, Bock R, Greiner S. 2017. GeSeq - versatile and accurate annotation of organelle genomes. Nucleic acids research 45: W6–W11.

UniProt Consortium. 2023. UniProt: the Universal Protein Knowledgebase in 2023. Nucleic acids research 51: D523–D531.

Werth S, Sork VL. 2014. Ecological specialization in Trebouxia (Trebouxiophyceae) photobionts of Ramalina menziesii (Ramalinaceae) across six range-covering ecoregions of western North America. American journal of botany 101: 1127–1140.

Wyczanska M, Wacker K, Dyer PS, Werth S. 2023. Local-scale panmixia in the lichenized fungus *Xanthoria parietina* contrasts with substantial genetic structure in its *Trebouxia* photobionts. *Lichenologist (London*, England): 1–11.

Xiao R, Zheng Y. 2016. Overview of microalgal extracellular polymeric substances (EPS) and their applications. Biotechnology advances 34: 1225–1244.

Yamamoto K, Hamaji T, Kawai-Toyooka H, Matsuzaki R, Takahashi F, Nishimura Y, Kawachi M, Noguchi H, Minakuchi Y, Umen JG, et al. 2021. Three genomes in the algal genus reveal the fate of a haploid sex-determining region after a transition to homothallism. Proceedings of the National Academy of Sciences of the United States of America 118.

Yin Y, Mao X, Yang J, Chen X, Mao F, Xu Y. 2012. dbCAN: a web resource for automated carbohydrate-active enzyme annotation. Nucleic acids research 40: W445–51.

Yoshimura I, Yamamoto Y, Nakano T, Finnie J. 2002. Isolation and Culture of Lichen Photobionts and Mycobionts. In: Protocols in Lichenology. Berlin, Heidelberg: Springer Berlin Heidelberg, 3–33.

Yu G, Wang L-G, Han Y, He Q-Y. 2012. clusterProfiler: an R package for comparing biological themes among gene clusters. Omics: a journal of integrative biology 16: 284–287.

